# Effect of short-term prescription opioids on DNA methylation of the *OPRM1* promoter

**DOI:** 10.1101/2020.01.24.919084

**Authors:** Jose Vladimir Sandoval-Sierra, Francisco I Salgado García, Jeffrey H Brooks, Karen J Derefinko, Khyobeni Mozhui

## Abstract

**Background:** Long-term opioid use has been associated with hypermethylation of the opioid receptor mu 1 (*OPRM1*) promoter. Very little is currently known about the early epigenetic response to therapeutic opioids. Here we examine whether we can detect DNA methylation changes associated with few days use of prescribed opioids. Genome-wide DNA methylation was assayed in a cohort of 33 opioid-naïve participants who underwent standard dental surgery followed by opioid self-administration. Saliva samples were collected before surgery (visit 1), and at two postsurgery visits at 2.7 ± 1.5 days (visit 2), and 39 ± 10 days (visit 3) after the discontinuation of opioid analgesics.

**Results:** The perioperative methylome underwent significant changes over the three visits that was primarily due to postoperative inflammatory response and cell heterogeneity. To specifically examine the effect of opioids, we started with a candidate gene approach and evaluated 10 CpGs located in the *OPRM1* promoter. There was significant cross-sectional variability in opioid use, and for participants who self-administered the prescribed drugs, the total dosage ranged from 5–210 morphine milligram equivalent (MME). Participants were categorized by cumulative dosage into three groups: <25 MME, 25–90 MME, ≥90 MME. Using mixed effects modeling, 4 CpGs had significant positive associations with opioid dose at 2-tailed p-value < 0.05, and overall, 9 of the 10 *OPRM1* promoter CpGs showed the predicted higher methylation in the higher dose groups relative to the lowest dose group. After adjustment for age, cellular heterogeneity, and past tobacco use, the promoter mean methylation also had positive associations with cumulative MME (regression coefficient = 0.0002, 1-tailed p-value = 0.02), and duration of opioid use (regression coefficient = 0.003, 1-tailed p-value = 0.001), but this effect was significant only for visit 3. A preliminary epigenome-wide association study identified a significant CpG in the promoter of the RAS-related signaling gene, *RASL10A*, that may be predictive of opioid dosage.

**Conclusion:** The present study provides evidence that the hypermethylation of the *OPRM1* promoter is in response to opioid use, and that epigenetic differences in *OPRM1* and other sites are associated with short-term use of therapeutic opioids.

## Background

Prescription opioids were once considered as a relatively benign treatment for pain management [1, 2]. However, over the past decade, prescribed analgesics have emerged as a major socio-environmental factor that has contributed to the opioid epidemic [3, 4]. For many individuals who develop opioid use disorder (OUD), the initiation phase may begin with treatment for acute pain or minor surgery, with primary care physicians and dentists accounting for a large fraction of prescribed opioids [5–11]. Even short-term use (e.g., up to three days) is a risk factor for some individuals, and the risk for addiction increases proportionally with dosage and duration of use [8, 9, 12–15].

Drug addiction is a chronic disease that is triggered by an exposure to an environmental agent. Following the initial exposure, the addictive substance continues to have a persistent effect, and this suggests a form of cellular memory. There is strong evidence that epigenetic processes, including DNA methylation, play a key role in maintaining the long-term effects of the additive substance [16, 17]. Studies particularly in model organisms have shown that drugs of abuse trigger intracellular signaling cascades that alter gene transcription; repeated exposure to the drug then results in remodeling of the epigenome that persists over time; and these epigenetic processes maintain the long-term changes in steady-state gene expression that underlie addiction [16, 18–20]. Work in humans generally relies on postmortem tissue from long-term drug users, and studies have found significant epigenetic differences in brains of former addicts compared to non-addicts [21, 22]. While the brain is the most relevant tissue in terms of neuroadaptation and drug seeking behavior, epigenetic markers of addiction have also been detected in peripheral tissues such as blood and sperm [23–28]. Easily accessible peripheral tissues are clearly the practical choice when it comes to defining biomarkers of drug use and/or predictors of individual risk for addiction.

The μ-opioid receptor gene (*OPRM1*) encodes the primary target for both endogenous and exogeneous opioids and plays a central role in mediating the rewarding and therapeutic effects. The CpG island located in the promoter of this gene is a potential sensor for drug use, and multiple studies in leukocytes and sperm have found higher DNA methylation among long-term opioid users compared to control samples [23, 29–33]. Hypermethylation of the promoter region has also been found among people with alcohol dependence [34]. However, as all these studies are cross-sectional comparisons between opioid-exposed individuals and controls, there is no definite way to discern whether the epigenetic differences are the cause, or effect, of drug use. Since genetic variants both within, and near the *OPRM1* gene have also been associated with susceptibility to addiction and drug sensitivity[35–37], it is plausible that such epigenetic markers represent genetic effects that preceded drug use. Another lingering question is, if the epigenetic changes are induced by drug use, does the hypermethylation of the promoter CpGs occur only after repeated and sustained exposure, or are these indicators of the early epigenomic, and potentially transcriptomic, responses to drugs? In the case of potent drugs such as opioids, the initial exposure is a crucial phase in the pathway to drug dependence and addiction, and it is reasonable to expect that some of the modification to the epigenome occurs within the first few exposures.

To address these questions, we applied a longitudinal design and collected saliva samples and self-reports of opioid use from a group of opioid naïve dental patients before oral surgery, and at two follow-up visits after surgery. We assayed genome-wide DNA methylation and explored (1) the methylome during the perioperative period, (2) how demographic variables such as age and race/ethnicity relate to methylome changes and immune response, and (3) whether we can discern opioid associated CpGs from the highly heterogeneous methylome data. As the site of surgery and postsurgery inflammatory response, the saliva presents particular challenges due to immune-related cellular heterogeneity. To overcome this, we applied *in-silico* approaches to deconvolute the underlying cellular heterogeneity and demonstrate the utility of the methylome-based cell estimates as proxies for the immune changes induced by surgery. For the effect of opioids, we specifically focused on the *OPRM1* promoter CpGs and evaluated whether the data replicates the CpG hypermethylation. Overall our results show a dose-dependent increase in methylation at the *OPRM1* promoter that can be discerned despite extensive heterogeneity in the methylome data, and this indicates that the epigenetic response to opioids occurs within the first few days to weeks following drug exposure. Additionally, we also performed an epigenome-wide association study (EWAS), and this identified a few CpGs that may be predictive of opioid dosing.

## Results

The number of enrolled participants (N = 41) and timeline of sample collection are shown in Fig. 1. Only 33 patients (19 females) received prescription opioids after an oral procedure. The baseline characteristics, other diagnosed diseases, casual use of other drugs (specifically tobacco and marijuana; no participant reported use of cocaine, psychedelics, and other hard drugs), and prescription opioid self-administration are reported only for these 33 participants (Table 1). Following the pre-surgery visit (visit 1 or v1), the second visit (visit 2 or v2) occurred after surgery and within a week of the last opioid dose (average number of days between last opioid dose and visit 2 was 2.7 ± 1.5 days). The last sample collection (visit 3 or v3) occurred between 32–88 days from surgery, and the number of days between the last opioid dose and visit 3 was 39 ± 10 days. In total, 26 participants provided saliva samples at all three visits, 6 participants provided saliva at two visits, and one provided saliva only at v1 (Table 1). The mean age was 33.61 ± 13.84 years and ranged from 19 to 61 years (Table 1). Based on self-reported race/ethnicity, there were 13 Caucasians (mean age = 31.69 ± 14.11 years), 13 African Americans (mean age = 39.92 ± 13.91), and the remaining 7 were of “other” racial/ethnic group (mostly Hispanic/Latino; mean age = 25.43 ± 8.02). The African American group was slightly older but there was no statistically significant difference in age between the groups (p-value = 0.06). Sex distribution was not significantly different between the race/ethnic groups. Individual level information, including comorbidities, is provided as Additional file 1: Table S1.

**Fig 1.**
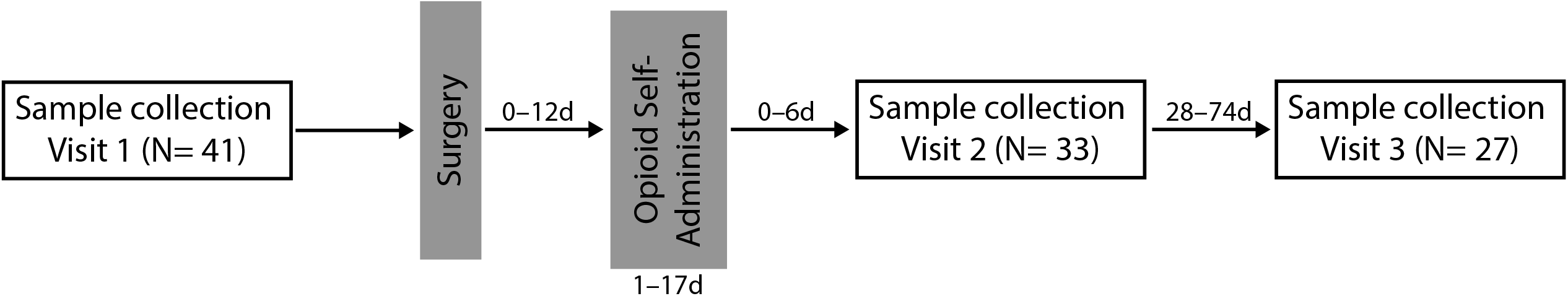
Timeline of sample collection. Saliva samples were collected before surgery and the two follow-up visits after surgery and end of opioid self-administration. The notations above the arrows show the range of days between events.

**Table 1.** Participant characteristics and postoperative opioid use.

| **Variables<sup>a</sup>** |  |
| --- | --- |
| **Sex** |  |
| Female | 19 |
| Male | 14 |
| **Age (years)** | $33.61 \pm 13.84$ |
| **Self-reported race/ethnicity** |  |
| African-American | 13 |
| Caucasian | 13 |
| Other<sup>b</sup> | 7 |
| **Tobacco in past 12 months<sup>c</sup>** |  |
| Yes | 11 |
| No | 22 |
| **Marijuana in past 12 months<sup>c</sup>** |  |
| Yes | 10 |
| No | 23 |
| **Other disease diagnosis<sup>c</sup>** |  |
| Yes | 17 |
| No | 16 |
| **Prescribed opioid medication** |  |
| Hydrocodone 5mg | 19 |
| Oxycodone 5mg | 12 |
| Oxycodone 10mg | 1 |
| Oxycodone 5mg and Codeine 30mg | 1 |
| **Length of opioid use in days (mean ± sd)** | $6 \pm 4$ |
| **MME<sup>d</sup> (mean ± sd)** | $64.91 \pm 48.88$ |
| <25 MME (<Q1) | 10 |
| 25–90 MME (Q1–Q3) | 13 |
| $\geq 90$ MME (≥Q3) | 10 |
| **Surgery to visit 2 in days (mean ± sd)** | $8.0 \pm 4.1$ |
| **Surgery to visit 3 in days (mean ± sd)** | $43.9 \pm 10.9$ |
| **Number of completed visits** |  |
| Three visits (v1, v2, and v3) | 26 participants |
| Two visits (v1 and v2) | 5 participants <sup>e</sup> |
| Two visits (v2 and v3) | 1 participant |
| Only v1 | 1 participant |

<sup>a</sup> Mean and standard deviation (sd) for continuous variables and counts for categorical variables
<sup>b</sup> Other= Hispanic/Latino, Asian, Middle-eastern, and Native American
<sup>c</sup> Self-reported data on other drug use and diagnosis of other diseases (diseases listed in Additional file 1: Table S1)
<sup>d</sup> Opioid dose converted to morphine milligram equivalent (MME) according to medication type; Q1 is the first quartile (25%) and Q3 is the third quartile (75%)
<sup>e</sup> Methylome data for one participant with v1 and v2 samples were excluded during the methylome data check (see methods)

Postoperative opioid dosing data was based on self-reported pill counts converted to morphine milligram equivalent (MME). With the exception of one individual who used no opioids (and we considered this individual to represent a dose of 0 MME with 0 days of use), all patients started opioid treatment generally within 24 hours of surgery, and continued use for an average of 6 ± 4 days for up to 17 days (Additional file 1: Table S1). As expected, cumulative dosage correlated with length of use (r = 0.67, p-value < 0.0001). For the 32 participants that self-administered opioids, the total cumulative dosage over the course of treatment ranged from 5–210 MME. Based on the quantile distribution of the cumulative MME, participants were classified into three groups: <25 MME (those below the 25^th^ percentile or quartile 1 for opioid dosage), 25-90 MME (those within the interquartile range), and ≥90 MME (those above the 75^th^ percentile or quartile 3) (Table 1). Opioid dosage showed no significant association with age, sex, and self-reported race/ethnicity. There was no significant association between opioid dosage and the presence or absence of other comorbidities. Dosage was also not associated with past marijuana use. However, the group that reported using tobacco within the past 12 months had significantly higher self-administered opioid dosage (mean of 95.63 ± 57.78 MME among tobacco users, and 49.55 ± 36.19 MME among non-users; p-value = 0.008).

### Global shift in postoperative methylome

For an overview of the methylome and the variance structure, we started with a principal component analysis (PCA) using the full set of high quality probes (736,432 probes passed QC criteria). The top PC (PC1) captured a vast portion of the variance at 63.5%, and following that, PC2 and PC3 captured only 2.5% and 1.6% of variance, respectively (PCs for each methylome data in Additional file 1: Table S2). PC1 was not significantly associated with the demographic variables (sex, age, self-reported race/ethnicity), or with comorbidities and past use of marijuana or tobacco. Instead, visit was the most significant explanatory variable for PC1 (F_2_,86= 5.94, p-value = 0.004), and the pattern indicated a significant change in the methylome with the strongest contrast between v3 and v2 (Tukey-Kramer *post hoc* p-value = 0.003) (Fig. 2a). To deduce whether the longitudinal variance capture by PC1 could be explained by the length of time from surgery or opioid self-administration, we performed bivariate analyses between PC1 and the following variables: opioid dose, days from surgery to sample collection, and days from last opioid self-administration to sample collection. This analysis was done for the three visits separately, and at v2, PC1 had a modest but significant correlation with days from surgery to v2 (r = 0.40, p-value = 0.03, n = 31 participants with methylome data at v2; Fig. 2b). Similarly, at v3, PC1 was correlated with days from surgery to v3 (r = 0.41, p-value = 0.04, n = 27 participants with methylome data at v3). PC1 was not correlated with opioid dose or the number of days from the last opioid use. From this, we can infer that the longitudinal shift in the methylome is primarily due to surgery.

**Figure 2.**
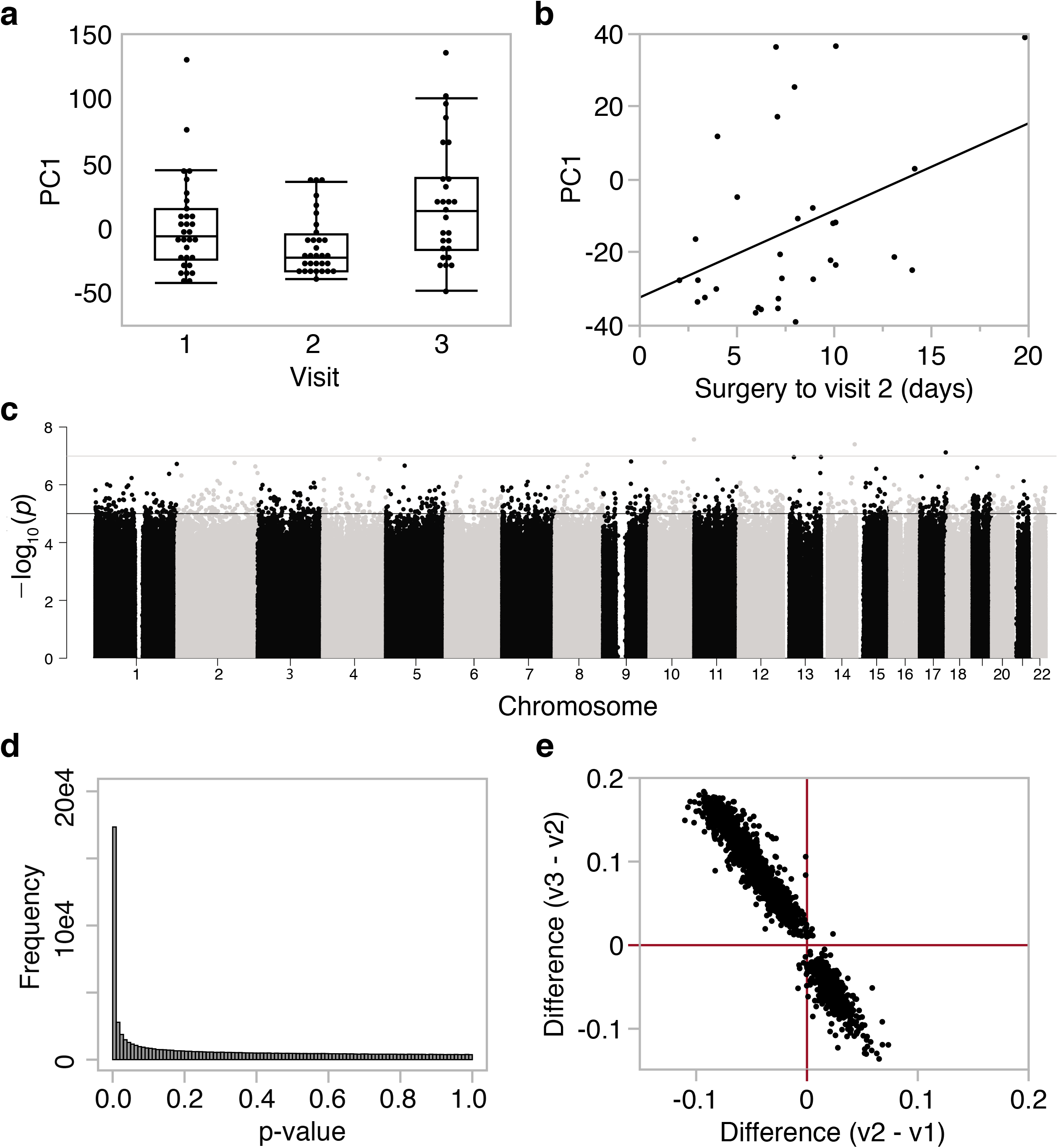
Global patterns in DNA methylation across visits. **(a)** The top principal component (PC1) extracted from the methylome-wide data explained 63.5% of the variance, and the ANOVA plot shows significant differences between the three visits (F_2_,86= 5.94, p-value = 0.004). (b) At visit 2, PC1 is correlated with number of days from surgery to the second visit (r = 0.40, p-value = 0.03, n = 31). (c) The epigenome-wide association plot depicts the location of each CpG (autosomal chromosomes 1 to 22 on the x-axis) and the – −log_10_(p-value) for the effect of visit (y-axis). Genome-wide significant threshold was set at p-value = 5 x 10^-8^ (upper red horizontal line); suggestive threshold was set at p-value = 10^-5^ (lower blue horizontal line), (d) Distribution of p-values for the effect of visit shows a significant deviation from the null hypothesis, (e) For the CpGs above the suggestive threshold, comparison of mean differences between visit 1 and visit 2 (x-axis), and visit 3 and visit 2 (y-axis) indicates a reversal in methylation patterns from visit 2 to visit 3, with the majority of sites showing lower methylation at visit 2, and then increasing in methylation by visit 3.

To profile the CpGs that changed longitudinally over the three visits we performed a mixed effects ANOVA with visit as a fixed variable and the person ID as random effect (Fig. 2c). The p-values for visit showed a significant deviation from the null hypothesis (Fig. 2d histogram). However, only 2 intergenic CpGs (cg05639411 and cg24904009) were above the genome-wide significant threshold of 5.0e-8 (Fig. 2c) and overall, the pattern indicated a modest shift in the methylome across several CpGs. At a genome-wide suggestive threshold of p-value = 1.5e-5, there were 1701 CpGs that underwent change over the visits (Additional file 1: Table S3). The majority of these CpGs (>65%) decreased in methylation between v1 and v2, and regained methylation by v3 such that these sites showed significantly higher levels of methylation at v3 compared to both v1 and v2 (Fig. 2e). Similarly, for the ~35% of CpGs that gained methylation between v1 and v2, these sites generally declined in methylation by v3 resulting in significantly lower methylation compared to both v1 and v2 (Fig. 2e). Gene set enrichment analysis (GSEA) of the 1133 annotated genes represented by the CpGs conveyed mostly an innate immune inflammatory response (Additional file 1: Table S4). The most overrepresented pathway was natural killer cell mediated cytotoxicity (KEGG ID hsa04650; normalized enrichment score = - 1.93, FDR = 0.03), and the most overrepresented function was for genes involved in cellular defense response (GO ID 0006968; normalized enrichment score = −1.83 p = 0.001, FDR = 0.3), and these immunity-related categories were enriched among the CpGs that decreased in methylation at v2. The opioid receptors were not represented in the list of visit associated CpGs. Based on these observations, a possible explanation for the shift in the methylome is that it is the result of surgery-induced immune response and changes in the oral cell composition. Opioid use, if it had an impact, is likely to exert a weaker signal, and given the limited sample size,more suitable for a focused candidate gene study.

### Deconvolution of cellular heterogeneity

To decompose cell types from the composite DNA methylation signal, we applied a reference-free approach [38]. The bootstrapping method described in Houseman et al. [38] determined K = 4 cell types (Additional file 1: Table S2). Cell 1, which represented the most abundant cell type, showed an increase at v2 right after surgery followed by a decline by v3 (Table 2). Aside from cell 1, no other cell showed significant change over the visits (Table 2). To deduce what cell types are represented by the 4 groups, we also estimated blood leukocyte proportions (mainly lymphocytes and granulocytes/neutrophils) using a reference-based approach [39], and compared correlations between the 4 cell types to the reference-based cell estimates (Table 2; Additional file 1: Table S2). Cell 1 had a strong positive correlation with granulocytes, and cell 4 had a strong positive correlation with lymphocytes indicating that cells 1 and 4 are chiefly representative of the leukocyte population in saliva, and serves as a proxy for the increase in granulocyte proportions after surgery. Cells 2 and 3 had only modest correlations with leukocyte estimates (|r| of 0.4–0.5) and may be more representative of the epithelial cells.

**Table 2.** Reference-free and reference-based estimates of cellular proportions.

| Cell types | Cell proportions by visit (mean ± SD) | | | | Pearson r with reference-based estimates | |
| --- | --- | --- | --- | --- | --- | --- |
|  | Visit 1 | Visit 2 | Visit 3 | Visit p-val | Lymphocytes | Granulocytes |
|  | *Reference-free estimates* |  |  |  |  |  |
| Cell 1 | 0.60 ± 0.28 | 0.74 ± 0.23 | 0.47 ± 0.32 | $F_{2,86} = 6.6$,<br>0.002 | -0.96 | 0.95 |
| Cell 2 | 0.13 ± 0.13 | 0.10 ± 0.10 | 0.17 ± 0.15 | ns | 0.51 | -0.47 |
| Cell 3 | 0.21 ± 0.20 | 0.12 ± 0.18 | 0.22 ± 0.19 | ns | 0.43 | -0.40 |
| Cell 4 | 0.07 ± 0.18 | 0.03 ± 0.08 | 0.14 ± 0.23 | $F_{2,86} = 2.7$,<br>0.07 | 0.77 | -0.81 |
|  | *Reference-based estimates* |  |  |  |  |  |
| Granulocytes | 0.71 ± 0.14 | 0.77 ± 0.10 | 0.64 ± 0.18 | $F_{2,86} = 6.1$,<br>0.003 | | |
| Lymphocytes | 0.27 ± 0.12 | 0.21 ± 0.09 | 0.33 ± 0.15 | $F_{2,86} = 6.6$,<br>0.002 | | |

The cell estimates were not associated with opioid dose. To evaluate if the baseline characteristics were related to cellular composition, we tested associations with age, sex, and race/ethnicity. Cell 1 had a significant negative correlation with age (r = −0.48, p-value = 0.006) only at v2 that suggests an age-dependent immune response in the days immediately after surgery (Fig. 3a). Cell 3 had the strongest association with age at all three visits (Fig. 3b). Cells 2 and 3 showed extensive cross-sectional variability without longitudinal change, and both were significantly associated with race/ethnicity at all three visits, indicating that these could serve as proxies for the cellular composition differences between populations (Fig. 3c, 3d). Cell 4 was not associated with any of the baseline variables, and sex was not a factor for any of the cell types.

**Figure 3.**
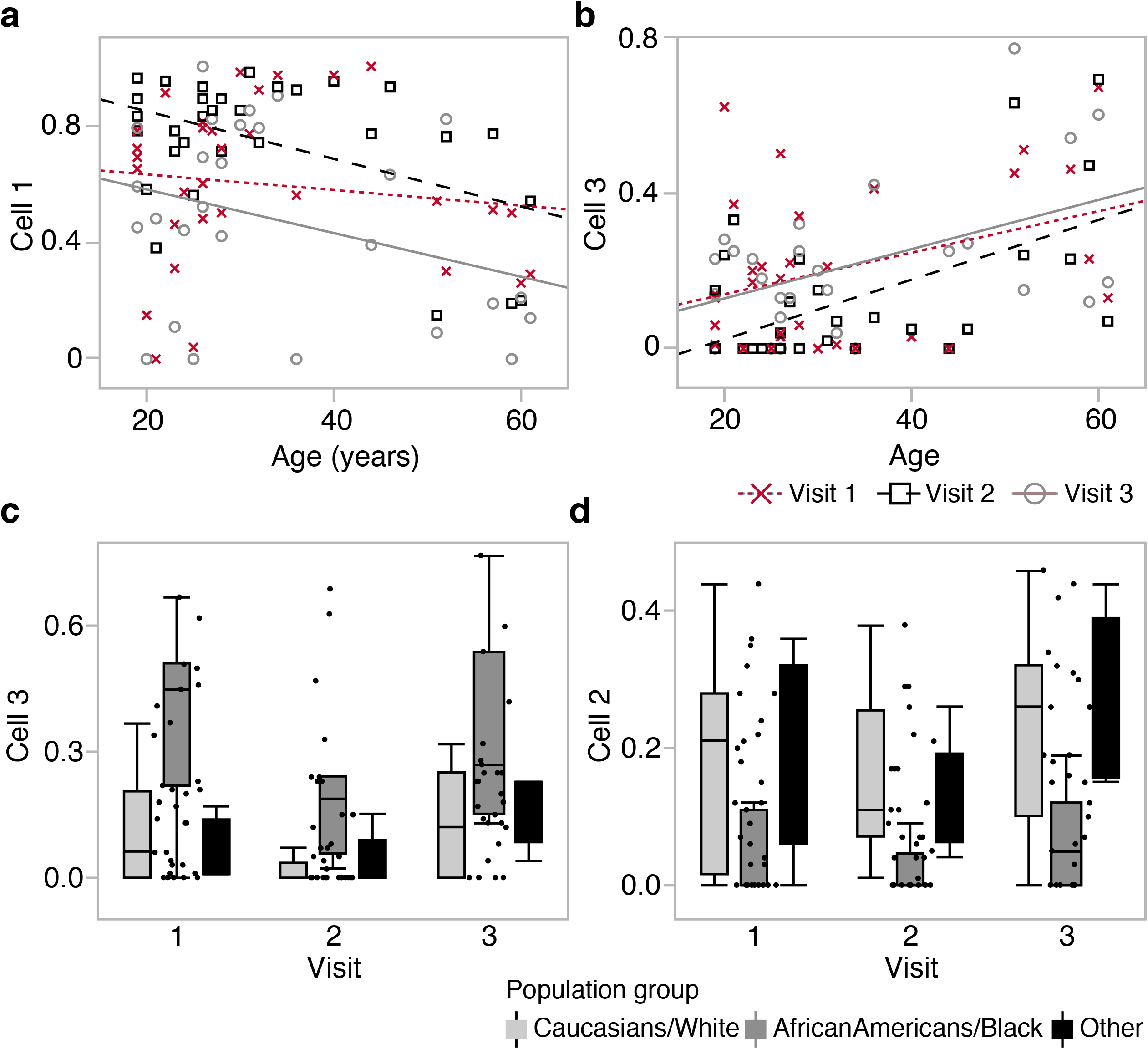
Estimated cell type proportions and associated variables. **(a)** Cell 1 shows both longitudinal and cross-sectional variability. Proportion of cell 1 is negatively correlated with age at visit 2 (r = −0.48, p-value = 0.006, n = 31; black squares and dashed line), but not at visit 1 (r = −0.13, p-value = 0.49, n = 31; red x markers and dotted line), and only slightly at visit 3 (r = −0.32, p-value = 0.10, n = 27; grey circles and solid line). (b) Cell 3 is associated with cross-sectional variability but no significant longitudinal change. The estimated proportion has a strong positive correlation with age at all three visits. At visit 1, r = 0.36 (p-value = 0.05); visit 2, r = 0.57 (p-value = 0.0009); visit 3, r = 0.48 (p-value = 0.01). (c) Cell 3 also shows a significantly higher proportion in African Americans at all three visits (F_2_,28 = 15.66, p-value < 0.0001 at visit 1). (d) Cell 2 is also ethnicity specific and associated with lower proportion in African Americans at all three visits (F_2,28_ = 4.77, p-value < 0.02 at visit 1).

### Effect of opioid dose on *OPRM1* promoter methylation

To examine if higher opioid dose is related to higher promoter methylation, we started with a candidate gene approach and focused on the CpGs located in the *OPRM1* promoter. In total, 10 promoter CpGs were targeted by the Illumina probes and these encompassed the CpG island described by Nielsen et al. and replicated by Chorbov et al. [29, 30] (Fig. 4a; Table 3; individual level β-values in Additional file 1: Table S2). We first applied a mixed regression model with opioid dosage group and visit as fixed categorical variables, and each participant ID as random intercept. With the exception of the last CpG, the regression estimates for all the *OPRM1* promoter CpGs were positive, with higher methylation levels for the two higher dosage groups (i.e., 25-90 MME and ≥ 90 MME) relative to the lowest dosage group (<25 MME) (Table 3). At a nominal p-value of 0.05, 4 CpGs were significantly associated with opioid dosage groups. The ANOVA plots for these CpGs showed that the difference between dosage groups was pronounced at v3 (for CpG1, CpG2, CpG6) and v2 (for CpG7) but not at v1 (Fig. 4a). As tobacco use was associated with higher self-administered dosage of opioids, we considered it as a potential contributing factor. However, including past tobacco use in the regression model did not alter the results, and this indicated that the higher methylation at the *OPRM1* CpGs is a specific effect of opioids.

**Figure 4.**
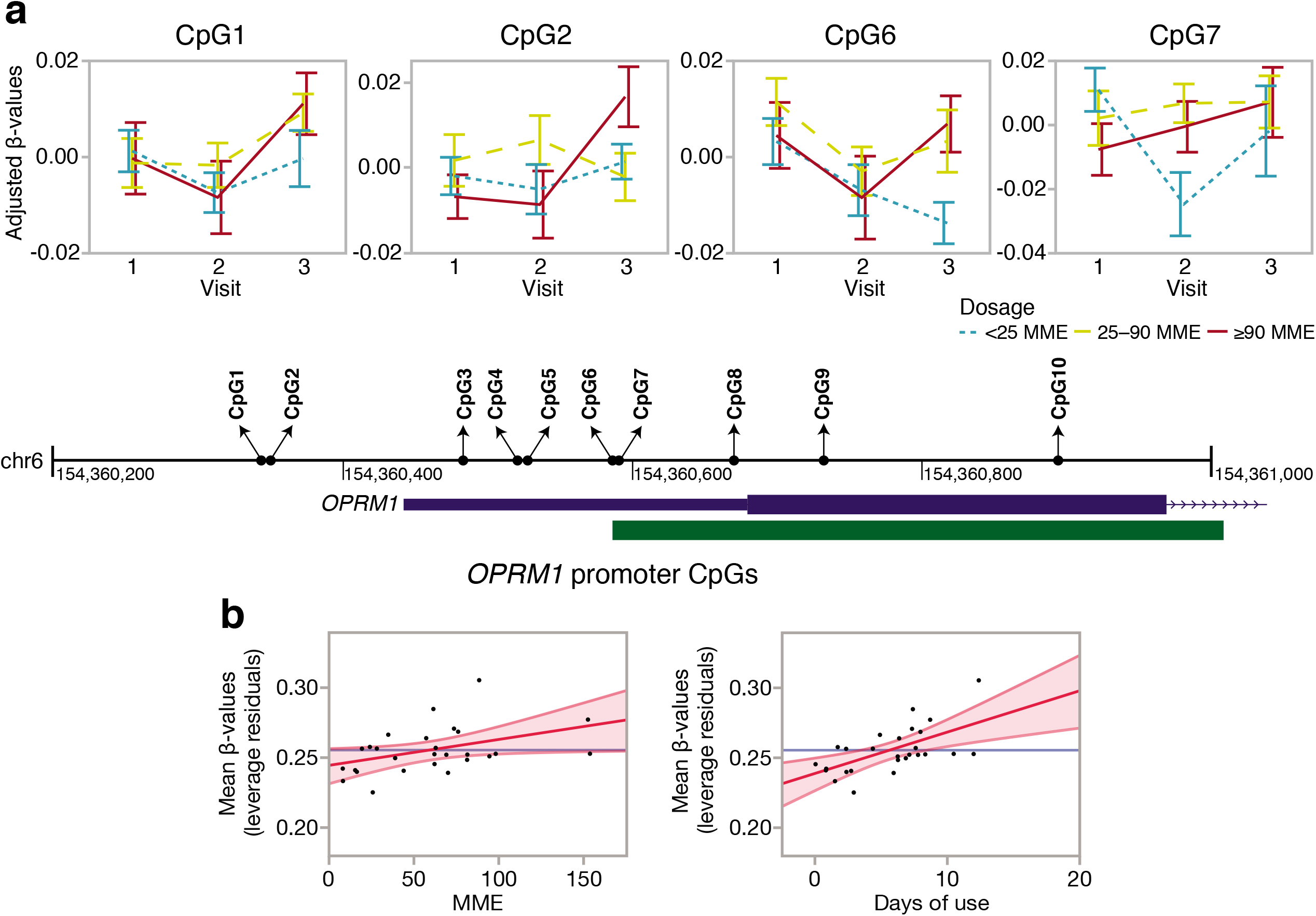
*OPRM1* promoter CpG methylation. **(a)** The *OPRM1* promoter and the CpG island (green block) are depicted along with base pair coordinates (black line; GRCh37/hg19), and location of the 10 CpGs (filled circles). Residual β-values were extracted after fitting participant ID as random intercept, and the plots show the methylation patterns across the three visits for CpG1, CpG2, CpG6, and CpG7 (panels with ANOVA line plots; error bars are standard error). The difference between the dosage groups appears at visit 3 (for CpG1, CpG2, CpG6) and visit 2 (for CpG7). The lowest cumulative dose group (<25 MME: blue dotted line) has lower average methylation compared to the two higher cumulative dose groups (25–90 MME: yellow dashed line; ≥90 MME: red solid line). (b) The promoter mean methylation was taken as the average β-values for CpG1 to CpG9. After fitting a regression model with adjustment for age and cell proportions, the leverage plots show a significantly higher average promoter methylation (y-axes) associated with higher MME (x-axis, left panel), and longer duration of use (x-axis; right panel). MME is morphine milligram equivalent.

**Table 3.** Dose dependent methylation of individual *OPRM1* promoter CpGs.

| CpG | ProbeID | <25 MME vs. 25–90 MME<sup>1</sup> |  | <25 MME vs. ≥90 MME<sup>1</sup> |  | Dosage anova<sup>2</sup> |  |
| --- | --- | --- | --- | --- | --- | --- | --- |
|  |  | Coef | t-val | Coef | t-val | F<sub>2,29</sub> | p |
| CpG1 | cg22370006 | 0.041 | 2.66 | 0.023 | 1.39 | 3.53 | **0.04** |
| CpG2 | cg14262937 | 0.051 | 2.77 | 0.014 | 0.69 | 4.20 | **0.02** |
| CpG3 | cg06649410 | 0.047 | 1.65 | 0.010 | 0.31 | 1.56 | 0.23 |
| CpG4 | cg23143142 | 0.018 | 1.56 | 0.000 | -0.04 | 1.71 | 0.20 |
| CpG5 | cg23706388 | 0.010 | 0.76 | 0.006 | 0.38 | 0.29 | 0.75 |
| CpG6 | cg05215925 | 0.019 | 2.73 | 0.012 | 1.59 | 3.74 | **0.04** |
| CpG7 | cg14348757 | 0.042 | 2.78 | 0.019 | 1.16 | 3.92 | **0.03** |
| CpG8 | cg12838303 | 0.026 | 2.08 | 0.022 | 1.64 | 2.38 | 0.11 |
| CpG9 | cg22719623 | 0.006 | 0.52 | 0.004 | 0.33 | 0.14 | 0.87 |
| CpG10 | cg15085086 | -0.029 | -0.94 | -0.040 | -1.19 | 0.78 | 0.47 |

<sup>1</sup>Regression estimates for higher dose groups (25–90 MME and ≥90 MME) relative to lowest dose group (<25 MME) based on linear mixed effects model: lmer(CpG ~ dose + visit + (1|ID))
<sup>2</sup>Two-tailed p-values for the main effect of dosage groups

To check whether the association with opioid dosage can be robustly detected, we summarized the overall methylation pattern in the promoter by taking the mean DNA methylation β-values for the nine CpGs that were positively associated with opioid dosage (CpG1 to CpG9). We applied a linear regression model and tested whether higher mean methylation was associated with either higher MME or longer length of opioid use. This analysis was done for the three visits separately and adjusted for age, tobacco use, and cellular heterogeneity. Both MME and length of opioid use were associated with higher mean methylation, but this effect was significant only at v3, further indicating that the hypermethylation of the *OPRM1* receptor is more likely a response rather than a predisposing factor (Fig. 4b; Table 4). Our results are consistent with the opioid associated hypermethylation and indicates that even a relatively short-term opioid use may induce an increase in methylation that is proportional to dosage at the *OPRM1* promoter.

**Table 4.** Mean methylation in the *OPRM1* promoter and association with opioid dose and days of use.

|  | MME dosage effect Visit 3<sup>2</sup> |  |  | Days of opioid use effect Visit 3<sup>2</sup> |  |  |
| --- | --- | --- | --- | --- | --- | --- |
|  | Coef | t-val | p (1-tailed) | Coef | t-val | p (1-tailed) |
| Promoter methylation<sup>1</sup> | 0.0002 | 2.16 | 0.02 | 0.003 | 3.40 | 0.001 |

<sup>1</sup>*OPRM1* promoter methylation summarize by averaging the β-values for CpG1 to CpG9
<sup>2</sup>Linear regression at visit 3, one-tailed p-value to test hypermethylation with higher cumulative MME or longer duration of use

### Preliminary epigenome-wide association study for opioid dose

Since the candidate gene approach indicated that the short-term use of prescribed opioids can have an impact on CpG methylation, we expanded the analysis to an EWAS using the same mixed model to test association with opioid dosage. A CpG (cg08105965) located in the promoter CpG island of the GTPase signaling gene, *RASL10A* (RAS like family 10 member A), was genome-wide significant (p-value of 5.0e-8; Fig. 5a). Unlike the pattern for the *OPRM1* promoter, the lowest dose group had significantly lower methylation level at both visits 1 and 3, indicating that the difference preceded opioid use (Fig. 5b). In addition to the *RASL10A* promoter CpG, 5 other CpG sites were associated with opioid dosage at the suggestive threshold (p-value of 1.0e-5; Fig. 5c–g). For most of these, the methylation differences were apparent at v1 and preceded opioid use. For these top CpGs, adjusting for past tobacco use did not alter the results, and none of these sites were significantly associated with tobacco use. To explore if any of the CpGs that were above the suggestive threshold have been previously implicated in opioid use or dependence, we referred to recent human EWAS for opioid dependence [24], and methadone treatment dosage [28]. Based on comparison of probe IDs and genes, none of the CpGs we report here have been previously linked to opioid related traits.

**Figure 5.**
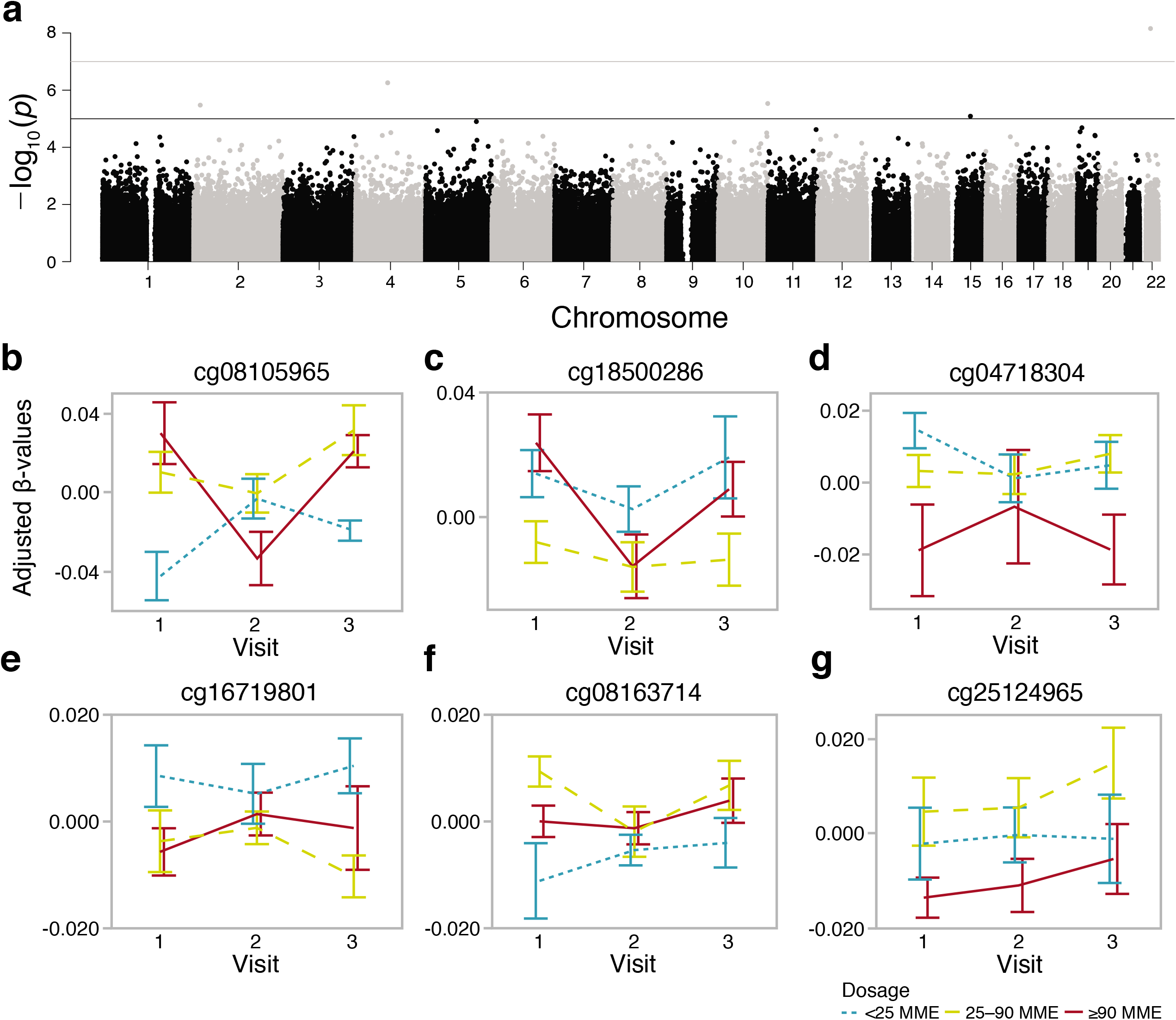
Epigenome-wide test for prescription opioid dosage. **(a)** The epigenome-wide Manhattan plot depicts the location of each CpG (autosomal chromosomes 1 to 22 on the x-axis) and the −log_10_(p-value) for the effect of doage (y-axis). Genome-wide significant threshold was set at p-value = 5 x 10^-8^ (upper red horizontal line); suggestive threshold was set at p-value = 10^-5^ (lower blue horizontal line). For the six CpG sites that were above the genome-wide suggestive threshold, residual β-values were extracted after fitting participant ID as random intercept, and the plots show the methylation patterns across the three visits (error bars are standard error) for **(b)** cg08105965 (*RASL10A*), **(c)** cg18500286 (*AFF1*), **(d)** cg04718304 (intergenic), **(e)** cg16719801 (*VSNL1*), **(f)** cg08163714 (*ANXA2*), and (g) cg25124965 (*PAIP2*). For most of these, the difference between the dosage groups appears at visits 1 and 3.

While this is preliminary results from a small study cohort and is yet to be replicated, we provide the list of 64 CpGs that were associated with opioid dosage at a nominal uncorrected p-value of 1.0e-4, along with the gene ontology IDs and KEGG pathways for the corresponding genes in Additional file 1: Table S5.

## Discussion

Here we report results from a longitudinal study of DNA methylation in a cohort of opioid naïve dental patients who received prescription opioids following oral surgery. To summarize the main result, we found increased methylation at the *OPRM1* promoter associated with higher cumulative opioid dose. This replicates the hypermethylated profile among long-term opioid users and alcohol dependent individuals [23, 29–34]. The pattern of methylation we observed indicates that the increase in methylation is more likely the response to, rather than the cause of, opioid use [20]. The present study provides evidence that such epigenetic modifications are induced within the early days of drug use and may represent early epigenomic responses to an addictive substance.

A peculiar challenge we faced was that the site of sample collection was also the site of surgery. Saliva has a highly heterogeneous cellular makeup and is estimated to constitute about ~45% epithelial cells, and about ~55% leukocytes from circulating blood [40]. The main goal of the study was to detect the effect of short-term and comparatively low-dose opioids, while accounting for the larger perturbation caused by surgery. Although we do not have details on the severity of the oral surgery, most were third molar extractions and were relatively minor and non-invasive. Nonetheless, the patients would have experienced an injury-induced inflammatory response that can result in changes in numbers of circulating immune cells [41], and consequently, changes in oral cell composition. As DNA methylation is highly cell-type specific, the heterogeneity in cells will be a major source of “noise” in the methylome data [39, 42–44]. The longitudinal variability in DNA methylation that was captured by the top PC can therefore be attributed to cell composition rather than opioid use. We could deduce this by the significant correlation between PC1 and the number of days from surgery to the follow-up visits. We were able to partly resolve the cell heterogeneity by applying reference-free and reference-based estimates of cell proportions. The reference-free method estimated four major cell types. Although saliva is highly heterogeneous, and certainly has more than just 4 types of cells [40], the classification into 4 broad groups likely reflects the limitation in the *in-silico* approach to resolve finer differences between cellular subtypes. Cell 1 most likely represented the granulocyte population (chiefly neutrophils), which constitutes the most abundant leukocyte subtype in circulating blood, and is responsible for innate immunity and acute inflammatory response. Consistent with the known increase in granulocyte-to-lymphocyte ratio in the few days following surgery [45], we also found an increase in cell 1 and in relative abundance of granulocytes compared to lymphocytes at visit 2. This was followed by a compensatory decrease in granulocyte proportions by visit 3. Cell 2 and cell 3 are presumed to represent a portion of the epithelial cell population, and these showed no significant within-individual changes over the visits. However, these cells exhibited significant association with age and self-reported race/ethnicity. Although cell type decomposition was not the primary objective of the study, our analyses demonstrated that the saliva methylome can be highly informative of individual differences in perioperative immune profiles.

For the effect of postsurgical opioid use, we first focused on the *OPRM1* promoter region as an epigenetic sensor of opioid dose. The CpG-rich promoter harbors a CpG island and several studies in different populations have demonstrated higher DNA methylation at this site among opioid users and methadone-maintained heroin addicts [23, 29–33]. The increased methylation of the *OPRM1* promoter is not only limited to OUD but has also been detected among individuals with alcohol dependence, suggesting that the hypermethylation is generally associated with substance use disorder and addiction [34]. A question has been whether such epigenetic differences are the result of drug use or the cause of increased vulnerability to addiction [20]. To address this, we interrogated 10 CpGs in a 550 bp region that encompassed the promoter CpG island investigated by Nielson et al. and Chorbov et al. (the CpG island is depicted in Fig. 4) [29, 30]. With the exception of the last CpG, the remaining 9 CpGs showed higher methylation in the two higher-dose groups relative to the low-dose group, and four of these CpGs were significantly associated with dosage at nominal alpha of 0.05. Comparison of mean methylation differences between the dosage groups across the three visits indicated that higher methylation in the higher dose groups is more apparent at the postsurgery visits, particularly visit 3. The positive association between the mean promoter methylation and cumulative MME, and mean promoter methylation and days of opioid use, were also significant only at visit 3. The heightened inflammatory state at visit 2, which occurred within a few days of surgery, may have been the reason why the more subtle effect of opioids was not significant at visit 2, and the positive association emerged only at visit 3.

The *OPRM1* locus presents a prime site for gene x environment interaction, a critical aspect of addiction since the addictive substance is an environmental agent that has a long-lasting biological effect. The *OPRM1* gene has been the subject of several candidate gene studies for addiction. Much attention has been paid to the missense SNP that alters the *OPRM1* protein function, although its impact on addiction traits and OUD is somewhat ambiguous [46, 47]. Several studies have also identified non-coding variants in the *OPRM1* locus that alters DNA methylation and gene expression [33, 35, 48]. At least one genome-wide association study has also identified a genome-wide significant association between a SNP upstream of *OPRM1* and methadone-maintenance dosing [37]. These studies collectively provide evidence that common genetic variants in the proximal region of *OPRM1* affect DNA methylation and gene expression, and could have a downstream impact on opioid response that could potentially influence vulnerability to addiction. Our present work was carried out in a small sample size and our primary goal was to track the within-individual trajectory across visits. If there were genetically modulated small cross-sectional differences at baseline, this sample size would be underpowered to detect the differences, and the significant association with opioid dose that we found may have been the result of opioid-induced augmentation of differential methylation at the postoperative visits.

The hypermethylation of the *OPRM1* promoter is likely only a small part of a larger network of genes involved in the cellular response to drug exposure. We therefore followed up with a preliminary EWAS exploration to identify other CpGs that may be associated with opioid self-administration. To our surprise, despite the small sample size, one CpG, located in the promoter region of *RASL10A*, a Ras-related GTPase signaling gene, was genome-wide significant. Perhaps this is due to the power of the longitudinal design in capturing differentially methylated sites that are significantly different between dosage groups at more than one visit. For instance, the differential methylation of cg08105965 at *RASL10A* is apparent at both visits 1 and 3. Although *RASL10A* has not been previously implicated in opioid response or addiction, it is notable that the OPRM1 protein is a G-protein coupled receptor, and its activation results in cellular signaling cascades that also involve Ras GTPase activity [49–51]. In addition to the *RASL10A* CpG, five other CpGs were at or above the suggestive threshold, including sites located in *AFF1, VSNL1, ANXA2*, and *PAIP2*. To our knowledge, DNA methylation at these genes have not been previously linked to opioid use or dependence. However, one recent study of gene expression in the rat model has shown an upregulation of *RASL10A* and *VSNL1* in the brain following acute morphine treatment [52]. Similar to *RASL10A, VSNL1* (visinin-like 1) also codes for an intracellular signaling molecule with high expression in the brain [53]. The list of CpGs that were associated with opioid dosage at a nominal uncorrected p-value < 0.0001 included few other cellular signaling genes (e.g., *ANXA2, RET, ADRB1*). Taken together, the EWAS results hint that epigenetic modulation of genes involved in intracellular signal transduction may play a role during the early phase of opioid use.

We must emphasize that the small sample size and the heterogeneity in methylome signal, partly due to cell composition and partly due to the heterogenous population group, are major limitations, and the EWAS results await replication in an independent cohort. The CpGs identified by the present EWAS were differentially methylated even at v1, prior to opioid use, and this suggests that there may be genetic variants underlying these epigenetic differences. However, such potential effects of genetic variation is not addressed in the present study due to the lack of genotype data. Another weakness that we should note is that the main variable of interest, therapeutic opioid dosing, was based on patient self-reports rather than objective measures of drug use [54]. A future strategy would be to use existing technologies such as wearable devices that can provide additional means of tracking the physiological responses to opioids [55]. The present study also does not address whether these epigenetic changes linger or diminish over time in the absence of continued drug use. A more comprehensive longitudinal epigenomic study of the early effects of prescription opioids that also integrates genetic effects, and with a longer follow-up period would be the next phase of study.

Regarding the potential for epigenetic persistence, we must point out that any peripheral tissue serves only as a proxy for the possible epigenetic changes in the brain. A distinction is that blood and epithelium are mitotically active tissues and cells are renewed within a few days to a few weeks, with the exception of long-lived memory T-cells. For methylation signals to persist, it will require either continued presence of the perturbation (i.e., continued exposure to opioids), or methylation changes in mitotically active stem cells that can be faithfully transmitted to daughter cells. The brain, on the other hand, is mitotically inactive and consists of mostly terminally differentiated cells that last a lifetime. If the relatively modest dose and short-term use of prescription opioids has a similar impact in brain cells, the effects may not readily decay and may be long lasting in the central nervous system.

## Conclusion

In conclusion, our study replicates the hypermethylation of the *OPRM1* promoter with opioid use. Previous studies reported on the effects of chronic opioid use; here we provide evidence that the epigenetic restructuring begins within the initial stage of opioid exposure. The present findings on the acute effects of prescription opioids, as well as the CpGs that may be predictive of opioid dosing, require further replication with a well-powered and more comprehensive study in a larger cohort.

## Methods

### Participants

Eligible participants were scheduled for tooth extractions, mostly third molar extractions, at an oral and maxillofacial surgery clinic that were typically followed by postoperative prescriptions of hydrocodone/acetaminophen (7.5/325 mg q4-6h prn pain) or oxycodone/acetaminophen (5mg/325mg q6h prn pain). For inclusion in the study, individuals were required to be 18 years of age or older, opioid naïve, able to consent, able to understand and speak English, and willing to provide saliva samples. Individuals were excluded if they reported previous use of opioids, had current substance use dependence, were pregnant, were incarcerated, had other causes of pain, were unable to consent, or had a developmental disability that prevented participation. The study received approval by the university Institutional Review Board. Eligible participants were provided a summary of the consent form by the study coordinator and allowed to read and ask questions before enrollment. All participants provided written informed consent.

Forty-one individuals consented to the study and provided contact information and responded to a demographic questionnaire. The enrolled participants also provided information on existing diagnosed diseases, and were assessed for casual substance use within the past 12 months (tobacco, marijuana, cocaine, psychedelics, other hard drugs; details in Additional file 1: Table S1). The clinical staff provided routine opioid medication and recovery instructions for all participants right after surgery. The opioids prescribed to participants were Hydrocodone, Oxycodone, and Codeine in doses that varied between 5mg to 30mg (Table 1). The study coordinator also provided opioid medication logs to participants to record self-administration including the date, time, individual dose per opioid pill, and number of opioid pills taken. For the 33 participants who received opioid medication, only one participant (person ID 142) reported no opioid usage.

### Sample processing and DNA methylation assay

Saliva was collected using the Oragene DNA sample collection kit (OGR 500) by DNA Genotek (http://www.dnagenotek.com). The first set of samples was collected before surgery. The second saliva sample was collected a few days after opioid discontinuation, and the third was collected on a follow-up visit (Fig. 1; Table 1). DNA was purified using the DNA Genotek PrepIT L2P kit according to manufacturer’s instruction. Genome-wide DNA methylation was assayed on the Illumina Infinium Human MethylationEPIC BeadChips following the manufacturer’s standard protocol at the HudsonAlpha Genomic Services Lab (https://gsl.hudsonalpha.org).

### Data processing

Raw intensity IDAT files were loaded to R and all quality checks, data preprocessing, and normalization were carried out using the R package, minfi (v.1.31) [56]. Methylation levels were estimated as β-values (ratio of methylated by unmethylated probes) and quantile normalized. The initial QC involved comparison between the log median intensities of methylated and unmethylated channels, and the density plots for β-values (Additional file 2: Fig. S1a). All samples passed these checks. Sex estimated from the DNA methylation data also matched the self-reported sex. To retain only high-quality data, probes with detection p-value > 0.01 (14,676 probes) were excluded. Probes that target CpGs on the sex chromosomes were also removed (18,605 probes). Finally, a total of 96,146 probes that overlapped annotated SNPs and/or were flagged for poor mapping quality (MASK.general list from [57]) were also filtered out. A total of 736,432 high quality probes were retained and used for downstream analysis.

As further QC, we performed unsupervised hierarchical clustering using the full set of high-quality probes (Additional file 2: Fig. S1b). While samples longitudinally collected from the same individual tended to cluster together, there were also several samples that did not cluster with self. To check for possible errors in sample labeling, we repeated the cluster analysis using a subset of 30,435 probes that had been filtered out due to overlap with common SNPs. While these were deemed poor-quality probes and unfit for differential methylation analysis, in terms of sample identity check, these probes can serve as proxy genotype markers that can help verify if samples came from the same person. Using this set, almost all samples collected from the same participant clustered with self, and for the most part, the clusters also aligned with self-reported race/ethnicity groups (Additional file 2: Fig. S1c). Only one of the 33 participant who received prescription opioids (person ID 108) did not pair with self and data from this person were excluded from all downstream analysis.

### Estimation of cellular proportions

To infer the relative proportions of the major cell types, we first implemented a reference-free deconvolution of the methylome data using the R package RefFreeEWAS (v2.2) [58]. The RefFreeEWAS algorithm applies a non-negative matrix factorization to decompose a matrix Y = MΩ, where M represents an *m x K* matrix with *m* as CpG specific methylation for an unknown number of *K* cell types, and Ω as the cell-type proportion constrained to sum to a value ≤ 1. For computational efficiency, the *K* cell types has to be first specified, and as described in Houseman et al. [58], we set the *K* to vary from 2 to 10 cell types and decomposed the Y = MΩ. Following this, we applied bootstrapping to estimate the optimal *K* value. For this estimation, we applied 10 iterations with replacement every 1000 times. The optimal *K* = 4 was selected based on the minimum value of the average of bootstrapped deviances for each putative cell type. While this method provides the relative proportions of cell types, the identity of the four cells are unknown. Since a significant proportion of saliva consists of leukocytes, we also applied a reference-based approach to estimate the relative proportions of lymphocytes and neutrophils [39, 42, 43]. To infer the putative identities of cells, we performed Pearson correlations between the *K* cells and the proportions of leukocyte types.

### Statistical analyses

For the global analysis, PCA was done on the full set of 736,432 probes using the prcomp function in R. In order to evaluate which variable had the most significant association with PC1, we examined the association between PC1 and the following variables: sex, age, self-reported race/ethnicity, presence or absence of other diseases, tobacco or marijuana usage in the past 12 months, and opioid dose, days from surgery to sample collection, and days from last opioid dose to sample collection. We used ANOVA for categorical variables, and Pearson correlation for continuous variables; and these tests were conducted separately for the three visits. We also performed similar analyses for estimated cell proportions to examine whether the variables were significantly related to the cell proportions. PC1 and the cell proportions were also related to visit using ANOVA. Since visit was the most significant explanatory variable for PC1, we identified the CpGs that showed longitudinal change over the three visits by applying a mixed-effects ANOVA: aov(β-value ~ visit + Error(ID/visit)). This epigenome-wide analysis was done for the 26 participants with data from all 3 visits. For the set of genome-wide suggestive CpGs that changed over the visits (uncorrected p-value ≤ 10^-5^), GSEA was implemented on the WebGestalt platform (http://www.webgestalt.org) with each CpG ranked by the mean β-value difference between v2 and v1.

For candidate gene analysis, we surveyed the promoter region of *OPRM1*. The CpG island that was interrogated by Nielson et al., and Chorbov et al. is located at 154360587–154360922 bp of chromosome 6 (GRCh37/hg19) [29, 30]. Within that exact coordinate, our data only had 4 CpG probes. We therefore considered a slightly wider region (550 bp) and in total, the array data contained 10 probes that targeted promoter CpGs at chr6:154360344-154360894 bp. To evaluate methylation at individual CpGs, we applied a linear mixed-effects model with dosage group and visit as fixed categorical variables, and person ID as random intercept: lmer(β-value ~ dosage + visit + (1| ID)). To test if history of tobacco use could account for some of the effects, we repeated the test with the model: lmer(β-value ~ MME + tobacco-use + visit + (1| ID)). This was done using the “Imertest” R package, and to get the p-values for the main effect of dosage groups, the degrees of freedom were computed by the Satterthwaite’s method [59, 60]. Following the CpG level analysis, we estimated the general methylation trend for the promoter by averaging the β-values for the 9 CpGs that had a positive regression coefficient with the dosage groups. We then tested the association between the promoter mean methylation score, and two opioid-related continuous variables: length of opioid use in days, and cumulative MME. This analysis was done for the three visits separately, and adjusted for age and cellular heterogeneity using the equations lm(mean-β ~ MME + tobacco-use + age + cell1 + cell2 + cell3), and lm(mean-β ~ days-of-use + tobacco-use + age + cell1 + cell2 + cell3).

Following the candidate gene study, we then performed an EWAS for opioid dosage using the same mixed-effect model: lmer(β-value ~ MME + visit + (1|ID)); and for the CpGs identified by the EWAS at above the genome-wide suggestive threshold, we checked for the effect of past tobacco use using the model: lmer(β-value ~ MME + tobacco-use + visit + (1|ID)).

## List of Abbreviations

EWAS: Epigenome-wide association study
FDR: False discovery rate
GO: Gene ontology
GSEA: Gene Set Enrichment Analysis
KEGG: Kyoto encyclopedia of genes and genomes
MME: Morphine milligram equivalent
OPRM1: Opioid receptor mu 1
OUD: Opioid use disorder
PC: Principal component
PCA: Principal component analysis
QC: Quality control
SNP: Single nucleotide polymorphism

## Declarations

### Ethics approval and consent to participate

All participants provided written informed consent and study received IRB approval.

### Consent for publication

Not applicable

### Availability of Data

The full de-identified raw DNA methylation data will be made available from the NCBI NIH Gene Expression Omnibus repository upon official publication.

### Author contributions

JVSS: performed lab work and data analysis and contributed to manuscript; FISG: contributed to data analysis; JHB: identified suitable patients and facilitated participant recruitment at the dental clinic; KJD: contributed to study conception and design; KM: contributed to study conception, design and data analysis, and wrote the manuscript. All authors contributed to and approved the final version of the manuscript.

## Acknowledgements

We thank the UTHSC Office of Research and the CORNET Awards for supporting this project.

