## Supplemental figure for "Effect of short-term prescription opioids on DNA methylation of the *OPRM1* promoter"

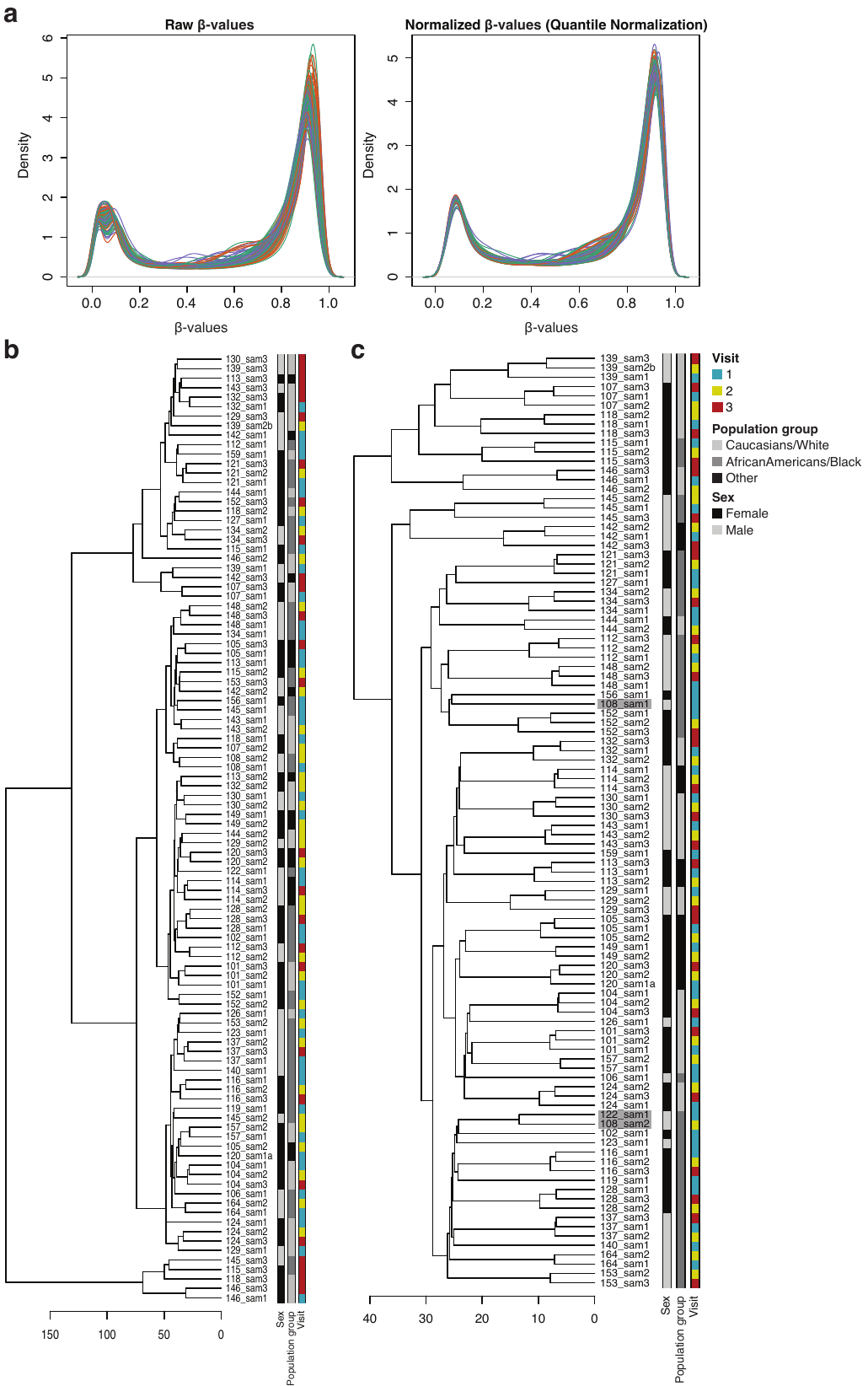

**Fig. S1.**  **Microarray data quality checks.** **(a)** The density plots for the β-values (left: raw values; right: quantile normalized) using the full set of >850K probes show the expected bimodal distribution. **(b)** Unsupervised hierarchical clustering using 736,432 high quality probes shows that while many of the longitudinally collected data cluster appropriately with self, several samples do not cluster with self. This may be due to within-individual heterogeneity over time; however, a concerning possibility is errors in sample identity. **(c)** Repeating the unsupervised hierarchical clustering using probes that were flagged due to overlap with SNPs shows that all samples collected longitudinally from the same participant cluster with self, with the exception of only one participant who received prescription opioids (greyed: person ID 108). Person 108 paired with a participant who received no prescription opioids (person ID 122) and had no follow-up data (both 108 and 122 were excluded from the main statistical analyses). Note that this QC checks were done using data from the full set of recruited participants; however, participants who received no prescription opioids (and therefore no follow-up visits) were excluded from the main analyses.
